# Whole genome sequencing of 56 *Mimulus* individuals illustrates population structure and local selection

**DOI:** 10.1101/031575

**Authors:** Joshua R. Puzey, John H. Willis, John K. Kelly

**Affiliations:** Department of Biology, College of William and Mary, Williamsburg, Virginia, 23187; Department of Biology, Duke University, Durham, North Carolina, 27708; Department of Ecology and Evolution, University of Kansas, Lawrence, Kansas, 27708

**Keywords:** population genomics, evolution, genomics, migration, selection, *Mimulus*, inversion

## Abstract

Across western North America, *Mimulus guttatus* exists as many local populations adapted to site-specific challenges including salt spray, temperature, water availability, and soil chemistry. Gene flow between locally adapted populations will effect genetic diversity in both local demes and across the larger meta-population. A single population of annual *M. guttatus* from Iron Mountain, Oregon (IM) has been extensively studied and we here building off this research by analyzing whole genome sequences from 34 inbred lines from IM in conjunction with sequences from 22 *Mimulus* individuals from across the geographic range. Three striking features of these data address hypotheses about migration and selection in a locally adapted population. First, we find very high intra-population polymorphism (synonymous π = 0.033). Variation outside genes may be even higher, but is difficult to estimate because excessive divergence affects read mapping. Second, IM exhibits a significantly positive genome-wide average for Tajima’s D. This indicates allele frequencies are typically more intermediate than expected from neutrality, opposite the pattern observed in other species. Third, IM exhibits a distinctive haplotype structure. There is a genome-wide excess of positive associations between minor alleles; consistent with an important effect of gene flow from nearby *Mimulus* populations. The combination of multiple data types, including a novel, tree-based analytic method and estimates for structural polymorphism (inversions) from previous genetic mapping studies, illustrates how the balance of strong local selection, limited dispersal, and meta-population dynamics manifests across the genome.

## INTRODUCTION

The *Mimulus guttatus* species complex is an enormous collection of localized populations, some of which are recognized as distinct taxa (e.g. *Mimulus nasutus*) [1, 2] or ecotypes (e.g. annual and perennial) [3]. Over the past two decades, *M. guttatus* has emerged as a powerful model system for addressing evolutionary and ecological questions related to local adaptation, inbreeding, and maintenance of variation [4, 5]. Endemic to western North America, *M. guttatus* has adapted to a wide range of habitats including serpentine barrens, metal-rich mine tailings, huge elevation ranges, and oceanic salt spray [6–9]. These populations are often inter-fertile to varying degrees. Gene flow occurs between populations [10], but potentially strong selection against immigrant genotypes [6] allows substantial genetic differentiation.

Previous studies have clearly shown the level of genetic differentiation increases with distance among populations in *M. guttatus.* Lowry and Willis [11] estimated F_ST_ = 0.48 across a set of 30 populations spanning a latitudinal range from 35–45 degrees. Similarly, Twyford and Friedman [12] obtained an F_ST_ of 0.46 for populations sampled across a large extent of *M. guttatus’* native range (latitudinal range: 31.2 – 53.8). F_ST_ is slightly lower (0.43) in a more geographically limited sampling of *M. guttatus* across Oregon (V. Koelling and J. K. Kelly, unpublished results). This survey included the Iron Mountain population (hereafter IM). If IM is compared to populations at much smaller distances (3 and 6 km, respectively), F_ST_ declines to 0.07 and 0.13, respectively [13]. Taken in total, these data indicate geographic limits on gene flow, suggesting that *M. guttatus* populations should be able to adapt to local environmental conditions. Local adaptation has been demonstrated in several reciprocal transplant experiments [6, 8]. When gene flow does occur, it is likely to introduce divergent alleles into a locally adapted population. Genomic migration-selection balance predicts that local populations experiencing high levels of gene flow will result in high genome-wide levels of nucleotide diversity [14, 15]. Genomic regions subject to local selection are expected to be less permeable to incoming haplotypes and should have lower nucleotide diversity, higher levels of absolute divergence between populations (Dxy), and distinct patterns of linkage disequilibria [14, 16–18].

To explore these predictions, we examine the population genomics of a single focal population in conjunction with data from the larger meta-population. Our focal population, IM, has been the subject of intense evolutionary and ecological research for the past 30 years. IM is an annual population where lifespan is strictly limited by water availability. During the short window between the spring snow melt and summer drought (routinely 6–10 weeks), seedlings must grow, flower, mate, and set seed. These abiotic pressures impose strong selection on IM [19, 20] and the population exhibits adaption to local conditions [6]. Despite this, IM retains high internal variability in both molecular and quantitative genetic traits [21, 22]. Recent studies have also revealed extensive structural (chromosomal) variation. Major inversions segregate both within IM (on Linkage Groups 6 and 11 [23, 24]) and also between IM and other populations (on Linkage Groups 5, 8, and 10 [3, 12, 25]). Inversions are predicted to directly influence migration-selection balance, and may have extensive effects on molecular variation owing to recombination suppression.

We use a combination of analytical techniques to characterize patterns of polymorphism, divergence and haplotype structure. By reconstructing distance-based trees for thousands of intervals across the entire genome, we evaluate patterns of relatedness and the possibility of gene flow into IM. Two primary conclusions emerge. First, despite the fact that IM is a localized population composed of only a few hundred thousand individuals, synonymous site diversity is among the highest reported for any organism. Second, the varying structure of polymorphism within IM relative to divergence from other *M. guttatus* populations indicates an overall genomic pattern of successful gene flow of divergent immigrant genotypes into IM. However, many loci show patterns indicative of local adaptation, particularly at SNPs associated with chromosomal inversions.

## MATERIALS AND METHODS

***Plant samples***: Approximately 1200 independent lines of *M. guttatus,* each founded from the seed set of a separate field-collected plant sampled from the IM [26]. Each line was subsequently maintained in a greenhouse by single-seed descent (self-fertilization) for 5–13 generations. As expected, these lines are almost completely homozygous at microsatellite loci with different lines fixed for different alleles [7]. DNA from 39 of these lines was newly generated and subsequently combined with data from 9 previously sequenced IM lines ([27]; reads downloaded from the JGI Short Read Archive). Sequence data from 18 “outgroups” (other populations or species in the complex; Supplemental Table 1) data was downloaded from SRA [27]. For all samples (both IM and outgroups), average read depth after filtering (as determined from VCF file using vcftools -depth) ranged from 2.6–24.27 (mean=6.68; calculated for genotyped bases on LG1, no indels included, Supplemental Table 1). We also sequenced the perennial species, *Mimulus decorus,* hereafter called Iron Mtn. Perennial (IMP). This species occurs in close proximity to the annual IM population.

**Table 1.** Contrast of population genetic statistics between monophyletic and polyphyletic windows.

|  | Monophyletic | Polyphyletic | p-value |
| --- | --- | --- | --- |
| $\pi$ | 0.012 | 0.014 | <0.0001 |
| Var[$\pi$] | 0.000045 | 0.000089 | <0.0001 |
| r | 0.084 | 0.135 | <0.0001 |
| S | 453 | 541 | <0.0001 |
| Tajima's D | 0.135 | 0.206 | <0.0001 |
| Dxy | 0.049 | 0.036 | <0.0001 |

***DNA extraction, library preparation, and sequencing***: We collected and froze leaf tissue for DNA extraction. We extracted DNA from leaf tissue using the Epicentre Leaf MasterPure kit (Epicentre, USA). Libraries for Illumina sequencing were made using the Illumina Nextera DNA kit (Illumina, USA), which utilizes a transposon-based system to integrate within and tag DNA. Individual barcodes were added during library preparation to facilitate multiplexing. Libraries were pooled in equal molar amounts based on concentrations measured using the Qubit high-sensitivity DNA assay and insert size distributions obtained from a Agilent bioanalyzer (HS-DNA chip) (Agilent Technologies, USA). Up to 24 libraries were pooled in a single Illumina HiSeq 2500 Rapid-Run sequencing run generating 150-bp paired-end reads.

***Alignment, Genotype Calling, and Residual Heterozygosity:*** After sequencing, we demultiplexed reads into individual samples and mapped them independently. Reads were aligned to the unmasked *Mimulus guttatus* v2.0 reference genome (http://www.phytozome.net/) using bowtie2 [28]. Next, we converted SAM alignment files to binary format using samtools [29] and then processed alignments with Picardtools (http://broadinstitute.github.io/picard; Commands: FixMates, MarkDuplicates, and AddReadGroups). The Picard processing validated read pairing, removed duplicate reads, and added read groups for analysis in the Genome Analysis Toolkit (GATK) [30]. GATK UnifiedGenotyper was used to call genotypes (details in supplement). Genotype VCF files were converted to tabbed format using vcftools (vcf-to-tab) [31]. Details can be found in supplementary methods.

After the initial genotyping, we masked putative SNPs that were excessively heterozygous (in more than 25% of lines) as these are likely due to mis-mapped reads. This is likely, for example, in regions of the reference genome where paralogs are incorrectly collapsed into a single gene. We further suppressed entire genomic intervals where, across lines, the mean heterozygosity dividing the average expected heterozygosity (given by the Hardy-Weinberg proportions) exceeds 0.5. Finally, for each individual, we calculated the ratio of observed to expected heterozygosity within 500 SNP windows across the genome. Within each line, we called a region heterozygous if the average was elevated across 10 successive windows. We identified a total of 429 residually heterozygous regions across all lines/chromosomes. This corresponds to 1.29% of the sequence in total. The Mendelian prediction for residual heterozygosity with single seed descent is 1.56% after 6 generations and 0.78% after 7 generations. The size distribution of putative residual heterozygous regions is consistent with the number of generations of selfing, the size of the genome 450–500 mB, and map length (about 125 cM per chromosome) (Supplemental Figure 1).

**Figure 1.**
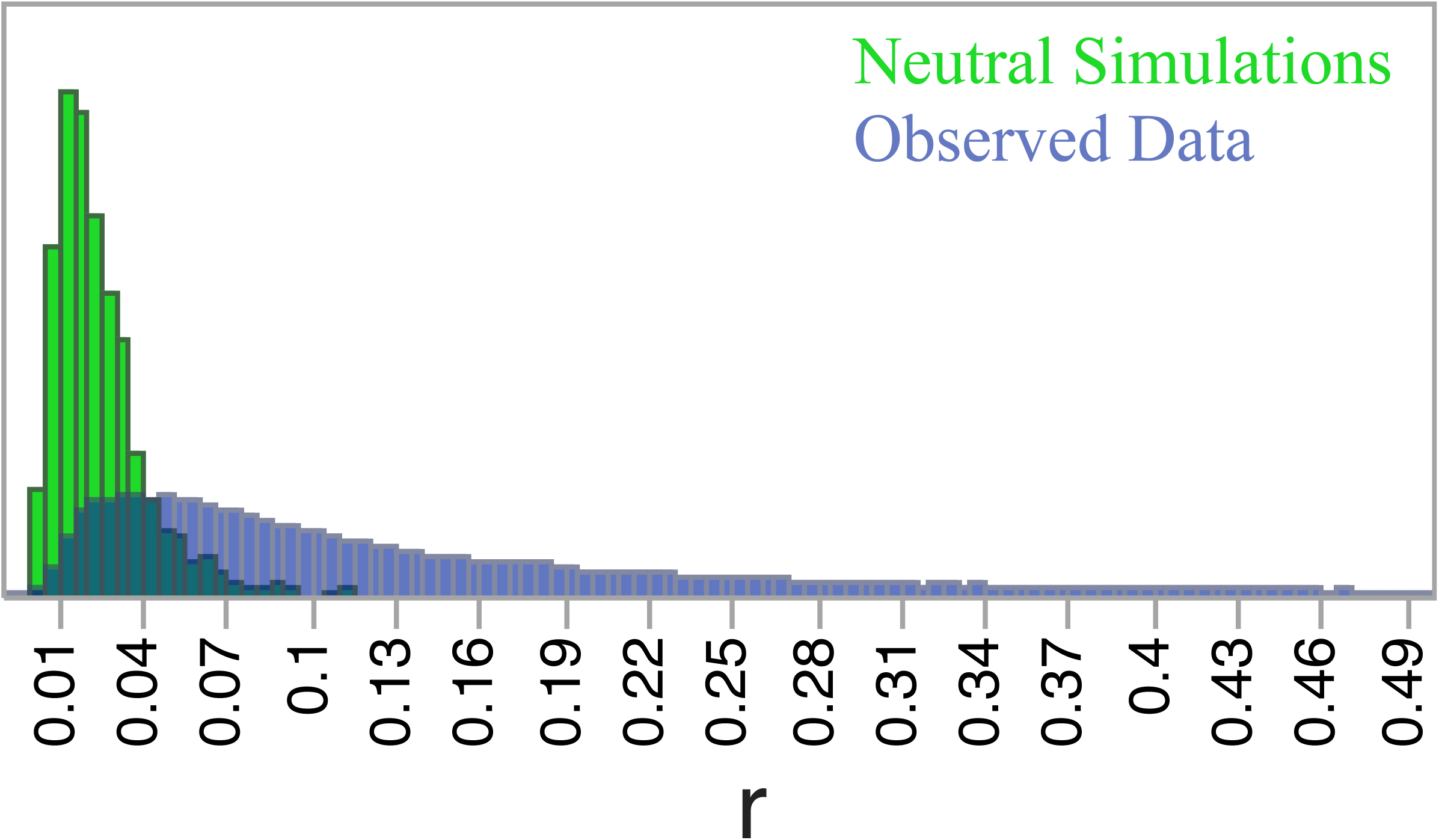
Genome-wide distribution of r in IM annuals shows an excess of positive associations between rare alleles relative to neutral simulations.

***Identification of related lines***: After the genotype filtering described above, we constructed a similarity matrix for all IM lines using the Emboss fdnadist program with the Jukes-Cantor substitution matrix. A total of 4.1 million SNPs were called in 43 or more IM lines. Based on this approach, we identified lines that were excessively similar (Supplemental Figure 2). For instance, IM777 is 0.997 similar to IM323. We thus determined these lines to be relatives and eliminated the IM323 from subsequent analyses. In addition, we also calculated the proportion of divergent sites through pairwise comparisons and used these values to identify related individuals. However, a low level of variability is consistent with a random sample from a large well-mixed population. IM109 is more divergent, but when considering the whole genome, is not a genuine outlier (Supplemental Figure 3). After filtering relatives, the following 34 lines were included for all subsequent tests: 62, 106, 109, 115, 116, 138, 170, 179, 238, 239, 266, 275, 359, 412, 479, 502, 549, 624, 657, 667, 693, 709, 742, 767, 777, 785, 835, 886, 909, 922, 1054, 1145, 1152, 1192.

***Relationship of missing data and divergence***: We delineated windows containing 500 SNPs, and within each window of each line, calculated (1) the number of called and uncalled sites and (2) the number of SNPs called for the reference allele as opposed to the alternative allele. The fraction missing data was calculated from (1), and the window divergence (fraction of calls to alternate) from (2). We performed a logistic regression in R with fraction missing as the response and divergence as the predictor: glm(formula = logit1$frac.missing ~ logit1$divergence, family = binomial). This revealed a strong relationship between data missingness and divergence (fraction of called SNPs that differ from the reference genome; Supplemental Figure 4A). Given this relationship, we opted to focus our analyses where data was most complete using two complimentary approaches. First, based on the fact that the fraction of missing data is considerably lower in coding regions (Supplemental Figure 4B) we conducted a series of gene-based analyses (e.g. synonymous versus non-synonymous diversity). The mean fraction of called bases in coding regions, calculated for each line, ranged from 0.63 to 0.86 (Supplemental Table 2). Second, we identified genomic windows (genic and inter-genic DNA) each consisting of 10,000 genotyped bases (monomorphic and polymorphic sites both count as genotyped bases). To qualify as a genotyped base, a site had to be scored in 30 of the 34 unrelated IM lines. The resulting windows ranged from 10,000–3,427,432 bases with a mean and median of 39,044 and 18,478 bases, respectively. Allowing 1,000 genotyped base overlapping steps between windows, a total of 74,445 windows span the 14 chromosomes. For these windows, we calculated population genetic and tree-based statistics.

***Nucleotide diversity within genes* (*Synonymous and non-synonymous π*)**: We converted filtered genotype files to fasta format for the entire genome for each separate line. When recreating line specific fasta files, missing data was not imputed, and indels and heterozygous sites were suppressed. Gffread [28] was used to extract coding sequences from individual fasta file. Each gene was individually extracted from the line specific coding sequences libraries and combined into a single fasta file containing 34 individual coding sequences for each gene. We calculated synonymous and non-synonymous diversity through pairwise comparisons of all lines using the KaKs_Calculator (Nei and Gojobori model) described in Zhang et al [32]. Only diversity measurements derived from genes with alignment lengths greater than 1000 bases were included (N=29,421). A Ka and Ks value was computed for each gene and was used to calculate genome-wide mean Ka and Ks.

***Window analyses***: We calculated statistics of polymorphism, divergence, and genealogy within windows of 10,000 genotyped bases. We created a phylogenetic tree for every window using EMBOSS fdnadist [33] to calculate a nucleotide distance matrix (Jukes-Cantor substitution model). Trees were inferred from the distance matrix using EMBOSS fneighbor [33] and rooted using *Mimulus dentilobius.* Of the 74,445 windows, fneighbor failed to parse the distance matrix for 28 windows. These windows were excluded from population genetic statistics results. Next, for each tree we determined whether IM was monophyletic or polyphyletic using a custom perl script [34] dependent on the Bio::Phylo toolkit [35]. This perl script searches a newick file and asks whether a specified group of individuals form a monophyletic clade. In the cases that IM was polyphyletic, we further explored the data by asking how many outgroup samples had to be removed to restore IM monophyly using the perl monophyletic output [34] and custom perl scripts.

We calculated S (the number of polymorphisms), π (nucleotide diversity), Tajima’s D [36], haplotype homozygosity, and LD statistics in each window using custom python scripts and VariScan [37]. For the linkage disequilibrium (D), we estimated the association of the minor alleles (less common base) at each contrasted SNP pair. Positive D indicates that minor alleles are positively associated [38]. We standardize D as the correlation coefficient, r = D/ Sqrt[p(1-p)q(1-q)], and from that, calculate r^2^ (the Z_nS_ test for selection is r^2^ conditioned on S [39]). We also calculated r and r^2^ for SNP pairs across each chromosome to estimate the long-range pattern of LD. For comparison to observed LD, we performed neutral simulations using calibrated, empirical estimates for 4Nu (from nucleotide diversity) and 4Nr (from LDhelmet as described below) by updating the programs used in Storz et al [40]. Absolute nucleotide divergence, Dxy, between IM annuals and all outgroups was calculated using a perl script [41] dependent on BioPerl::PopGen modules (Dxy is equivalent to π_XY_ [42]). Dxy was calculated on a single base increment for all sites that had at least one IM and one outgroup individual genotyped. Next, using these values, an average Dxy value was calculated for the same 10,000 genotyped base windows used for other population genetic statistics.

***Recombination rates within IM***: We used LDhelmet [43] to estimate fine-scale recombination rates with recalled genomes in fasta format as inputs. First, using the “find_confs” command, 50 SNP windows were used to scan the genome and create a haplotype configuration file. Next, a likelihood lookup table and Pade coefficients were generated using a population scaled mutation rate of 0.015 (this was based on a preliminary estimate for genome-wide π within IM). In the final step, the “rjmcmc” command was run using the previously generated haplotype configuration, likelihood table, and Pade coefficients and a Jukes-Cantor mutation matrix to estimate recombination rates. Exon specific recombination rates were calculated. Using bedtools [44] intersect command, coordinates of exons extracted from the *M. guttatus* gff3 gene annotation file (phytozome.net) were combined with recombination rates calculated by LDhelmet [43]. Only pairs of SNPs contained within exons were used. Using this information, we are able to look at gene and exon specific recombination rates.

## RESULTS

***Nucleotide diversity within IM***: Within genes, synonymous nucleotide diversity is very high π_syn_ = 0.033 (Supplement Figure 5). Mean non-synonymous diversity is π_non-syn_ = 0.006 (Supplemental Figure 5). Nucleotide diversity was significantly lower in interior exons (Supplemental Figure 6). Exon numbers one through five had mean π values of 0.0122, 0.0111, 0.0102, 0.0098, and 0.0092, respectively. Exon specific GC content was correlated with nucleotide diversity; π and GC content were both highest in the first exon (Supplemental Figure 6). When comparing the first exon to numbers 2–5, interior exons have lower GC (%GC_exon#1_=48.0, %GC_exon#2–5_=44.4, p<0.0001, *t-test*). To explore this relationship further, exons were placed in bins based on position (1^st^, 2^nd^, 3^rd^ exon, etc.) and further binned by GC content (40–45, 45–50, 50–55, and 55–60% GC content). Interestingly, in the first exon, π exhibits a negative relationship with GC content, while in exons 2–5, π is positively related with GC content (Supplemental Figure 7). Within the 10,000 genotyped base windows of IM lines (genic and inter-genic DNA), the average nucleotide diversity was 0.014. Across the genome, π varied from 0.00–0.03 (Supplemental Figure 8A and 9).

***Linkage disequilibrium and recombination rate:*** In most genomic windows there is a striking excess of positive LD (Figure 1). A large proportion of genomic windows exhibit stronger association of minor alleles than predicted with neutrality. If an IM line harbors the less frequent base at a SNP, it is much more likely to have the less frequent base at neighboring SNPs. The neutral distribution of Figure 1 was obtained using the average, genome-wide ρ (4Nr) of 0.0042 obtained from LDhelmet [43]. When measured as r^2^, linkage disequilibrium is high at short distances (~100bp) and shows a rapid decay with sequence distance (Supplemental Figure 12). It should be noted that the pattern of long-range LD differs among chromosomes (Supplemental Figure 12). The average genic ρ was 0.0052 and recombination hotspots were clearly evident (Supplemental Figure 13). On an exon specific level, recombination rates are highest in the first exons and decrease in interior exons (Supplemental Figure 6) (r_exon#1_=0.0073, SE=0.0002; r_exon#2–5_=0.0037, SE=0.0002; p<0.0001, *t-test*). Interestingly, in the first exon, π exhibits a negative relationship with GC content, while in exons 2–5, π is positively related with GC content (Supplemental Figure 7).

***Divergence of IM lines from other M. guttatus populations***: Diversity within IM was lower than divergence of IM sequences from outgroups (Dxy): Mean Dxy = 0.038, range 0.002–0.166 (Supplemental Figure 8E) with several clear peaks of high Dxy (Figure 2). The variance in pairwise divergence (among the 561 contrasts between 34 lines within each genomic window) also exhibits many localized peaks across the genome (Supplemental Figure 10). The mean of Var[π] is 0.0000831, and this statistic is positively correlated with nucleotide diversity in the window and with LD measured as r or r^2^ (Supplemental Figures 8F and 10, 11).

**Figure 2.**
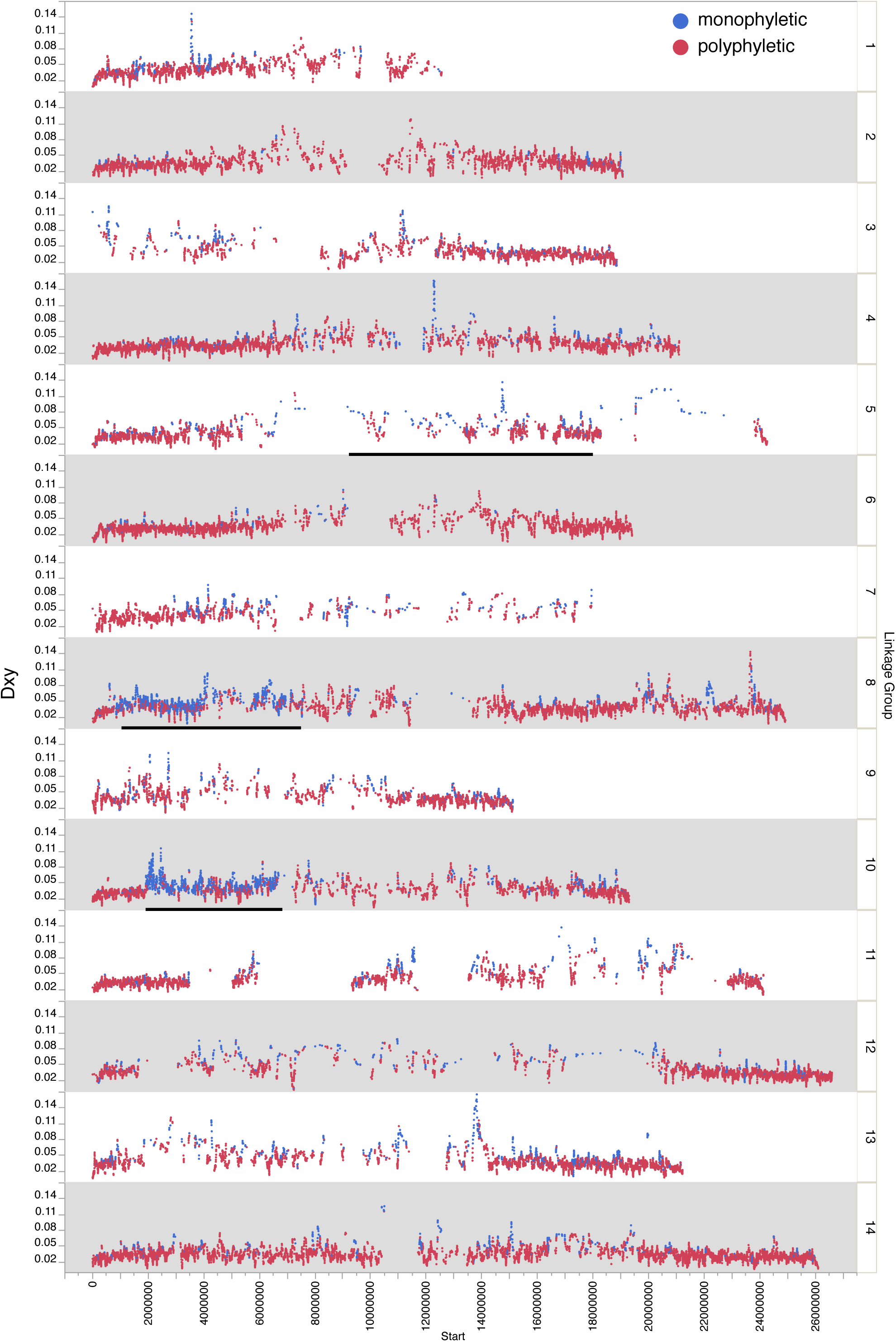
Genomic distribution of absolute divergence (Dxy). Blue dots = monophyletic windows; red dots = polyphyletic windows. Bars on LG5, LG8, and LG10 denote approximate locations of chromosomal inversions.

We located each of the three inversions mapped in the IMxPR RIL population [25] by locating the markers to locations in the v2 genome build (bars in Figure 2). This is a cross between annual (IM) and perennial (PR) genotypes. Absolute sequence divergence (Dxy) is significantly elevated within all three inversion regions relative to genome-wide averages: Dxy_Genome_ = 0.037, Dxy_inversion(Lg8)_ = 0.044 (p<0.0001), Dxy_inversion(LG5)_ = 0.068 (p<0.0001), and Dxy_invemon(LG10)_ = 0.042 (p<0.0001). The LG10 and LG8 inversions shows significantly lower overall nucleotide diversity while the LG5 is on not statistically different from genome-wide levels: π_Genome_=0.014, π_inversion(LG10)_ = 0.010 (p<0.0001), π_inversion(LG8)_ = 0.012 (p<0.0001), and π_inversion(LG5)_ = 0.014 (p=0.6603). Interestingly, Var[π] is statistically elevated in the LG5 inversion but statistically lower in the LG10 and LG8 inversions: Var[π]_Genome_= 0.000085, Var[π]_inversion(LG5)_ = 000109 (p=0.0002), Var[π]π_inversion(LG10)_ =0.000060 (p<0.0001), and Var[π]π_inversion(LG8)_ =0.000053 (p<0.0001).

***Distribution of monophyletic IM clusters across the genome:*** We constructed a phylogenetic tree for every window in the genome including both IM and outgroup samples. The monophyly of IM samples was evaluated individually for each tree. In 10,504 phylogenetic trees all IM samples were monophyletic (Supplemental Table 4). The majority (64,913) of windows showed IM as polyphyletic – some IM lines more similar to outgroup sequences than other IM lines (Supplemental Table 4). Genome-wide, IMmonphyletic windows were found both in gene dense and gene sparse regions. To further explore this data, we calculated the number of outgroup samples that would have to be removed from the tree for IM to be monphyletic. For 6,958 of the trees, only one outgroup sequence would have to be removed to restore IM monophyly (Supplemental Table 5). For 48% of these 6,958 trees, the geographically proximate Iron Mtn. Perennial (IMP) was responsible for IM paraphyly (Supplemental Table 5). This pattern is evident across all linkage groups but the incidence is highest for LG8 where 65% of polyphyletic trees where IM monophyly is ruined by a single individual are due to IMP (Supplemental Table 5).

***Relationship of Var[π] and tree topology:*** Combining within population Var[π] and tree topology allows for identification of genomic regions that have experienced a selective sweep or introgression event (Figure 3 and Supplemental Figure 15). On average, we expect monophyletic regions of the genome to have lower Var[π]. Introgression of divergent haplotypes should elevate Var[π], a trend we observe (Table 1). To test the effectiveness of the Var[π] statistic as a technique for identifying selective sweeps, we selected the lowest Var[π] tree from all monophyletic windows and looked at its topology and distribution of pairwise π. The tree created from the lowest Var[π] window shows a topology indicative of a selective sweep – short branches within the monophyletic IM clade. For this same window, within IM pairwise-n values are very small supporting the finding that all IM individuals within this block possess nearly the exact same haplotype (Figure 3A).

**Figure 3.**
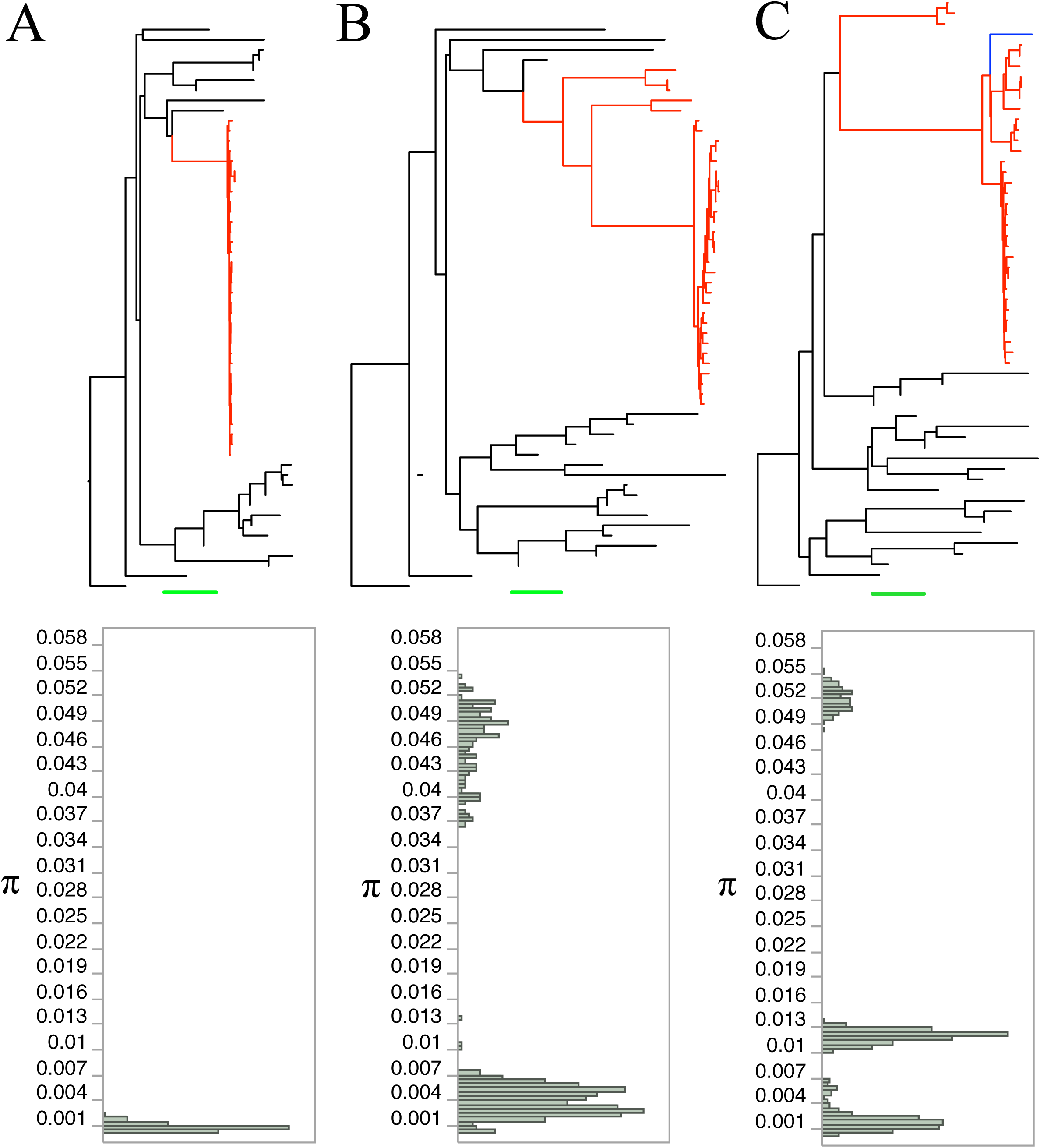
Relationship of tree topology and distribution of pairwise π values within IM annuals. Highlighted red branches are IM annuals. Black bars are outgroups. Green scale bar equals 0.01 for panels A, B and C. Pairwise π values between all 34 IM annuals were calculated (total 561 comparisons). Var[π] was calculated using these values. **(A)** Lowest Var[π] region for all monophyletic windows shows evidence of selective sweep. Top: tree shows that all IM annuals are very similar. Bottom: Distribution of raw π values for within IM comparisons shows that all IM annuals possess almost the exact same haplotype. **(B)** Highest Var[π] region for all monophyletic windows shows evidence of multiple distinct haplotypes within IM. **(C)** Second highest Var[π] region of all trees where IMP alone ruins IM monophyly shows evidence of introgression event including IMP and multiple distinct segregating haplotypes. Blue branch=IMP.

Next, we picked the highest Var[π] region for all monophyletic windows and created a tree and π distribution for this region (Figure 3B). These results indicate that for this genomic window the IM population contains several highly diverged sequences. Introgression may well have been the original source of the divergent lineages in Figure 3B, although local they may be maintained by selective processes within IM. It is also possible to use this dataset to identify the genomic effects of introgression from individual outgroup sequences. To illustrate this, we extracted all regions of the genome where IMP is solely responsible for breaking IM monophyly and extracted high Var[π] regions. Tree and π distributions from these windows show evidence of introgression and segregation of very divergent hapolotypes (long branches within IM and multimodal π distribution) (Figure 3C, Supplemental Figure 15).

***Contrast of population genetic statistics between monophyletic and polyphyletic regions:*** To determine the relationship between gene trees and population genetic statistics, we compared the distribution of each statistic between IM monophyletic windows and IM polyphyletic windows. Nucleotide diversity (π), Var[π], polarized LD (r), the number of segregating sites, and Tajima’s D were all significantly elevated in polyphyletic windows when compared to monophyletic windows (Table 1). Dxy was significantly lower in polyphyletic windows relative to monophyletic windows (Table 1).

## DISCUSSION

Traditionally, sequencing efforts in evolutionary genomics have focused on sampling a single individual from each of multiple populations distributed across the full range of a species. Only recently have evolutionary biologists begun generating whole-genome sequence datasets specific to demes, populations of individuals connected by mating in recent time, e.g. [45]. This study uses a combination of intensive within population and across population whole-genome sequencing to explore genomic patterns indicative of selection and introgression. In this discussion, we summarize genome-wide trends, discuss methodological approaches (tree based analysis combined with Var[π]) to explore within population sequencing data, and take a first step from genomic observations to underlying genes.

Nucleotide diversity within Iron Mountain (IM) is extremely high for a single population (π_Genome_ = 0.014, π_syn_ = 0.033). The genomic estimate of 0.014 is likely underestimated given the strong tendency for missing data to increases with divergence from the reference genome (Supplemental Figure 4). Downward bias of π outside of genic regions results from ascertainment; we are less likely to map (and thus analyze) sequences that are most divergent. This is not surprising, but to our knowledge, has not been clearly demonstrated previously. We expect this to be a general phenomenon extending across most studies of this kind. Here, we suspect that the high level of within population insertion/deletion variation reported for IM annuals contributes to incomplete mapping and subsequent underestimation of nucleotide diversity [27].

Leffler [46] has recently summarized nucleotide diversity across a wide-range of species, classifying estimates by sampling strategy (one population or multiple populations), site type (synonymous, non-synonymous, etc.), and chromosome type (autosome or sex). From their dataset, we extracted 37 one-population autosomal nucleotide diversity estimates spanning eight Phyla (Arthropoda, Chloropyta, Chordata, Echinodermata, Magnoliophyta, Mollusca, Pinophyta, and Porifera). Considering single population autosomal π_syn_ (N=9, species from five phylum represented: Arthropoda, Chlorophyta, Chordata, Mollusca, and Pinophyta), Leffler [46] observed a mean and median of 0.014 and 0.011, respectively (range 0.001–0.033). Across multiple populations (N=50) for which autosomal π_syn_ has been reported, values ranged from 0.000–0.035 with a mean and median of 0.010 and 0.006, respectively. It has previously been reported that *Mimulus guttatus* has multiple-population synonymous diversity values of π_syn_=0.061 [47]. Thus, for both single population and across multiple-populations, *Mimulus guttatus* ties and exceeds, respectively, the highest levels of autosomal synonymous nucleotide diversity reported. It is worth noting of all autosomal nucleotide diversity (π) values reported by Leffler (N=207) (not filtered based on site-type), four-fold degenerate site diversity (π=0.080) of the sea squirt (Ciona roulei) was the only higher π value than *M. guttatus* multiple population π_syn_=0.061 [46]. These results indicate that *Mimulus guttatus* has one of the highest within population and within species level of nucleotide diversity known for any organism.

Nucleotide diversity at a locus is determined by the number of SNPs and the frequencies of alternative alleles at each SNP. Relating to the latter, a second notable feature of IM is the substantially positive mean value for Tajima’s D (Table 1). Genomic surveys in several Drosophila species [45, 48, 49], as well as in Arabidopsis [50], reveal negative mean values for Tajima’s D. In other words, the allele frequency spectrum is skewed towards rare alleles in most genomic windows. In contrast, we find a tendency for allele frequencies to be more intermediate than expected from the equilibrium neutral model (predicted mean zero for Tajima’s D). The meiotic drive locus on chromosome 11 illustrates this pattern: Mean Tajima’s D = 0.46 across 561 genomic windows from position 9.5 Mb to 11.7 Mb. Previous study of this region has revealed a balanced polymorphism [24, 51]. An allele that causes meiotic drive has recently increased to a population frequency of approximately 35%. The genomic signature of this sort of recent event is subtle, unlike ancient balanced polymorphisms where alternative alleles may have highly divergent sequences [52]. In essence, the Drive haplotype has been ‘sampled’ from the SNPs resident in the population, and its increase owing to selection is associated with incremental shifts in the allele frequencies across a large genomic region. It is remarkable that the sequence-level pattern of variation within the Drive locus – a particular haplotype is present in a non-trivial minority of lines (20–30%) – is observed at many loci in the genome.

***Genome-wide effects of migration and localized signatures of selection***: Most genomic windows exhibit polyphyly and a haplotype structure suggesting low but significant migration of immigrant genotypes into IM. The former is simply the observation, that in most of the genome, some IM lines are more similar to lines from other populations than to other IM lines. The high frequency of polyphyly is not caused by a few divergent IM lines repeatedly breaking monophyly. Instead, most IM lines exhibit similarity to outgroup sequences within portions of their genomes; the identity of lines that break monophyly changing across the genome. This is expected given previous evidence that IM is an internally well-mixed, outbred population [53].

IM polyphyly could be due to either ancestral polymorphism that is continuing to segregate or introgression from neighboring populations. Both are likely relevant, but two observations suggest an important contribution of migration. First, we observed a specific pattern of LD in which the rarer alleles at pairs of SNPs are positively associated on average across the genome (Figure 1). This tendency is significantly elevated in polyphyletic windows (Table 1). Most sequencing studies do not specify associations between SNPs in terms of features of alternative alleles, and as a consequence, the direction of LD is meaningless. Indeed, direction is lost when calculating r^2^, which is commonly used to measure the strength of association between SNPs (e.g. Supplemental Figure 12). Here, in order to evaluate the potential contribution of introgression, we polarize alleles based on allele frequency in IM. Measured this way, the absolute value of LD can inform questions about evolutionary process. For example, Langley and Crow [38] developed an epistatic selection model to explain the negative LD observed in allozyme data. In IM, LD is highly variable in both direction and magnitude, but we suggest that the positive average is due, at least in part, to migration. Immigrants from divergent populations tend to generate positive LD by introducing novel alleles in combinations. The second observation is that the geographically proximate outgroup IMP is the most frequent cause of IM polyphyly (when a single outgroup is the cause; Supplemental Table 5). This suggests that migration is contributing to elevated π and Var[π] and that geographically proximate species are contributing divergent haplotypes.

Despite these trends, many genomic windows exhibit a distinct pattern suggesting local adaptation. The clearest signature of selection is a genomically localized reduction in nucleotide diversity coupled with increased divergence of IM from other populations of *M. guttatus* (Figure 2) [54]. One of the most striking observations from Table 1 is the elevated Dxy, depressed π and Var[π] in monophyletic windows. Reduced π is a one signal of directional selection while elevated Dxy is indicative of locally beneficial variants. Biologically, these statistics indicate the presence of locally favored variants present at high frequency with low levels of within population variation – all pointing to a local selective advantage.

The variance in pairwise nucleotide differences, Var[π], is an informative statistic, particularly when combined with gene-tree reconstructions (Table 1, Figure 3). Filtering genomic windows by Var[π] demonstrated that low Var[π] regions may have topologies indicative of selective sweeps (extremely short branch lengths), while high Var[π] regions exhibit multiple segregating haplotypes separated by longer branch lengths. These patterns suggest the genomic regions are subject to distinct evolutionary forces. For instance, balancing selection or soft sweeps may cause two haplotypes to be present at intermediate frequency within a population and would result in a local increase the variance of pairwise π. Another possible cause of elevated Var[π] would be sampling of a recently immigrated haplotype for which there has been insufficient time for recombination to breakdown. Patterns of LD may be used to distinguish between these two possibilities as large extended swaths of elevated Var[π] may be indicative of a recent introgression events, while sharp localized Var[π] peaks may be due to extended periods of balancing selection.

*Chromosomal Inversions:* We expect that selection effects will be most pronounced when recombination is suppressed. Recent studies suggest reduced gene flow between annual and perennial ‘ecotypes’ of *M. guttatus* within a chromosomal inversion on LG8 [3, 12]. Interestingly, QTL studies demonstrate that genes contained within this inversion are important for life history type and flowering time [3]. Holeski et al [25] recently mapped two additional putative inversions in a cross between IM and a perennial *M. guttatus* genotype from Point Reyes, CA. These are located on linkage groups 5 and 10 respectively (Figure 2). QTLs for traits related to reproductive isolation map to the LG8 inversion [3]; while the phenotypic effects of the other two loci remain to be investigated. The present dataset reveals that sequence divergence (Dxy) is significantly elevated within all three inversion regions relative to genome-wide observations (Figure 2). The LG10 and LG8 inversions shows significantly lower overall nucleotide diversity within IM, while the LG5 inversion is on not statistically different from genome-wide levels.

These results provide an interesting contrast to the literature on “genomic islands of speciation [55].” Genome scans in a number of systems have identified loci with elevated divergence among populations (populations sometimes described as species or nascent species) and these regions may harbor changes that reduce gene flow among populations. Cruickshank and Hahn recently reviewed the data from five systems (rabbits, mice, Heliconius butterflies, mosquitos, and flycatchers) and found that while relative divergence (measured by Fst) is higher in genomic islands, absolute divergence (measured by Dxy) is not (see Table 1 in [14]). They argue that this pattern, in conjunction with other observations, support the hypothesis that genomic islands are formed post-speciation. In other words, the relevant loci may not have actually reduced gene flow prior to speciation. The inversions in *M. guttatus* provide an interesting contrast to these examples for two reasons. First, they exhibit elevated divergence in both relative and absolute terms. Second, most of the outgroups included this survey (Supplemental Table 1) are described as divergent populations of one species (*M. guttatus*) rather than distinct/nascent species. Several self-fertilizing lineages (*M. nasutus, M. micranthus, M. platycalyx*) were included, but even for these, there is evidence of ongoing gene flow or introgression in the recent past [10, 56]. While gene flow seems to persist within this species complex, the populations that we contrast to IM are actually much more divergent (in terms of Dxy) than among taxa in four of the five systems reviewed by Cruickshank and Hahn (mice, butterflies, mosquitos, and flycatchers).

Patterns of genetic variation across the LG8 inversion within the IM population are surprising. First, the reduced intra-IM π for the LG8 inversion is noteworthy given that other studies have reported higher inter-population annual π than perennial π [57]. Second, it is very clear that the fraction of IM monophyletic trees is elevated within the LG8 inversion. The ratio of monophyletic to polyphyletic windows genome-wide (outside inversion) is 0.14 (9,049 monophyletic to 62,988 polyphyletic windows) while within the LG8 inversion this ratio is 1.58 (1455 monophyletic windows to 925 polyphyletic windows). Third, Var[π] is reduced and Dxy is elevated relative to genome-wide patterns. Fourth, of IMP is the sole cause of IM polyphyly far more frequently within the LG8 inversion than genome-wide.

Within the LG8 inversion, the fraction of IMP broken IM monophyly is considerably higher than genome-wide values. IMP is responsible for 409 of 925 (44%) polyphyletic windows, while outside the inversion, IMP is solely responsible for 2923 of 62,988 (4.6%) polyphyletic windows.

These data suggest that alleles contained with the LG8 inversion are important in local adaptation of the IM annuals. This is supported by the finding of reduced within IM π, reduced Var[π], dramatically increased frequency of IM monophyletic windows, and increased Dxy within this region. These data strongly suggest that and these regions are resistant to introgression due to selective pressures. Second, these data provide evidence for gene exchange via recombination or gene conversion between IM and IMP within LG8 region. This is supported by the fact that IMP is the sole cause of IM polyphyly far more frequently within the LG8 inversion that genome-wide. Put simply, IM and IMP share a unique set of sequences, not shared with outgroups, within the lg8 region more frequently than elsewhere in the genome. This will be of potential interest given the observation that the LG8 inversion is the genomic marker most closely association life-history of *M. guttatus* across its entire range.

Like the LG8 inversion, the LG10 inversions exhibits reduced intra-IM variation but increased divergence. In contrast, intra-IM π within the LG5 inversion is comparable to the genome-wide average. The very elevated Dxy, high Var[π], but moderate π within the LG5 inversion could be explained by reduced gene flow with outgroup populations and balancing local selection. The proportion of monophyletic windows within the inversion was substantially elevated when compared to outside LG5 inversion ratios. Within the LG5 inversion, 35 of 77 (45%) of windows were monophyletic. The proportion of monophyletic windows within the LG10 inversion is even more elevated (700 of 1330 (52%) windows monophyletic). Interestingly, IMP is not solely responsible for breaking IM monophyly in any of the 42 polyphyletic LG5 windows while IMP is the solely responsible for breaking IM monophyly in 241 of 630 LG10 windows.

The variable patterns of ancestry across inverted regions is perhaps not too surprising. Many different population/species of the M. guttatus complex are potential contributors to IM, and they may differ in the whether they have the IM orientation for a particular inversion.

*Candidate genes for future study: Mimulus guttutus* annuals have evolved a “live-fast-die-young” life history. To reproduce, they must take advantage of a narrow window of time between the spring snowmelt and summer drought. Abiotic tolerances and timing of life processes (germination, flowering, seed set, etc.) are particularly important in the success of IM annuals. Using the data assembled here, we sought to provide the foundation to move from overall genomic signals to identifying genes potentially important in these processes and possibly involved in local adaptation of the IM annual population. To begin to identify genes that are potential candidates of strong selection within the IM population, we calculated outlier residuals from a Dxy vs. π contrast (for all monophyletic window) and selected the 2.5% most extreme outlier windows where IM π is low for absolute divergence Dxy (Supplemental File 1; interval list of outlier windows). A total of 882 genes were located in these outlier windows, and they have significantly lower Ks (Ks_outlier_=0.014, Ks_Genome_=0.035, p<0.0001; for genes with alignment length >= 200 and p-value <=0.05, see methods) and lower non-synonymous diversity (Ka_outlier_=0.002, Ks_Genome_=0.006, p<0.0001). Genes involved in the flowering pathway, germination timing, stress responses, and trichome development are present in this outlier classes. While it is not possible to mention all interesting outlier genes, several very intriguing candidates are worth mentioning for follow-up functional work. *DELAY OF GERMINATION1* (DOG1), a gene involved in timing of germination and flowering in *Arabidopsis thaliana* [58, 59], as well as several genes involved in flowering time in *A. thaliana,* including *Short Vegetative Phase* (*SVP*) [60] and ATMBD9 [61], were contained within the outlier windows. The fact that these genes regulate phenological transitions in *A. thaliana* and that phenology is critical for IM fitness suggests that research following up on these candidate loci may move us a step closer to a mechanistic understanding of local adaptation.

## Data Availability

All sequence data generated here will be available on the Short Read Archive following acceptance of the manuscript. SRA numbers will be listed here following acceptance.

## Acknowledgements

We would like to thank Patrick Monnahan and Jenn Coughlan for providing extensive comments on this manuscript. This work was supported by grants from the National Institutes of Health to J.K. and J.W. (R01 GM073990) and the National Science Foundation to J.R.P. (NPGI-IOS-1202778).

## Author Contributions

JP, JW, and JK designed this experiment. JP made the libraries and directed sequencing. JP and JK performed all genomic analyses and wrote the paper.

